# Transcriptomic Insights into the Responses in Leaves Storing Lipid Organelles under Different Irradiances

**DOI:** 10.1101/2021.05.09.443314

**Authors:** Somrutai Winichayakul, Richard Macknight, Zac Beechey-Gradwell, Robyn Lee, Hong Xue, Tracey Crowther, Philip Anderson, Kim Richardson, Xiuying Zou, Dorothy Maher, Shona Brock, Luke Cooney, Gregory Bryan, Nick Roberts

**Affiliations:** Resilient Agriculture, AgResearch Ltd., Tennent Drive, Palmerston North 4442, New Zealand; Department of Biochemistry, University of Otago, Dunedin 9016, New Zealand

## Abstract

To increase the nutritional value of forage, transgenic ryegrass known as <u>H</u>igh <u>M</u>etabolizable <u>E</u>nergy (HME) were previously generated that co-express cysteine-oleosin and diacylglycerol O-acyltransferase. HME not only accumulate lipids in the leaf but also has elevated CO_2_ assimilation and increased biomass. Shading is one of the most influencing factors for ryegrass growth environments particularly in swards. The aim of this study, therefore, was to determine the influence of irradiance levels on photosynthesis and gene expression in the HME leaves when compared with their corresponding non-transformant (NT). Under low light (150-250 µmol m^-2^ s^-1^) and standard light (600-1000 µmol m^-2^ s^-1^), the HME accumulated more lipid than NT. The previously reported elevated photosynthesis and increased biomass was observed when the HME were grown under standard light but not under low light. Under both light conditions, compared to NT, the HME had upregulated a number of transcripts involved in lipid metabolism, light capturing, photosynthesis, and sugar signalling network while downregulated genes participated in sugar and fructan biosynthesis. We further discuss how the HME differentially manipulated several genes other metabolic pathways including maintenance of redox homeostasis. Combined, the data suggests that the increased photosynthesis capacity in the HME likely corresponds to an increase of micro-lipid sink strength; these are influenced by available light energy and may be related to diffusional and biochemical activities of stomata. Overall, this work provides a clearly understanding of the changes in molecular and biochemical mechanisms underlying the carbon storing as leaf lipid sink of the HME ryegrass.

**One sentence summary:** Shading led to increase leaf lipid accumulation but limit the greater photosynthesis trait of high lipid ryegrass which may be related to limitation of biochemical activities of stomata.

## INTRODUCTION

The climate crises and increasing world population means dramatic increases in the production of food, forage and renewable fuels is needed using the existing agricultural land. To address this, we and others have been exploring biotechnological approaches to increase the levels of energy-rich triacylglycerol (TAG) content in plant non-seed tissues (Xu and Shanklin 2016; Vanhercke et al., 2019, and references therein). Increased TAG synthesis in vegetative tissues can be achieved by promoting the carbon flux into fatty acid (FA) and TAG synthesis through the manipulation of enzymes and/or transcriptional regulators whilst suppressing FA catabolism, including FA utilization and TAG turnover. Co-expression of diacylglycerol O-acyltransferase (DGAT, the enzyme for TAG biosynthesis) and WRINKLED1 (WRI1, a key regulator of lipid homeostasis in the leaves) in a sugar-dependent TAG lipase (*sdp*1) mutant of tobacco yielded 5-8% dry weight (DW) TAG accumulation; however these plants incurred a growth penalty (Kelly et al., 2013). Other approaches include using a double mutant of *sdp1* and *trigalactosyldiacylglycerol 1-1* in Arabidopsis resulting in approximately 8% leaf TAG (%DW) while co-expressing DGAT1, WRI1, OLEOSIN and the lipid-related transcription factor LEAFY COTYLEDON 2 in tobacco leaves resulted to 23-30% TAG accumulation (Fan et al., 2014; Vanhercke et al., 2017, respectively). The tobacco plants also had a reduced biomass however; although this was reported to be partially offset by manipulating genes involved in carbohydrate metabolism (Mitchell et al., 2020). We previously reported that the development of transgenic lines of ryegrass (*Lolium perenne*) and Arabidopsis, known as high metabolizable energy (HME) plants, that co-express DGAT and cysteine-oleosin (cys-OLE, a modified oleosin designed to protect lipid droplets from proteolysis). These plants have up to twice the FA content in mature leaves (from ∼ 3.5% to ∼ 7% DW) as well as increasing plant CO_2_ assimilation rate and biomass (Roberts et al., 2010 & 2011; Winichayakul et al 2013 & 2020; Beechey-Gradwell et al., 2018 & 2020; Cooney et al., 2021). We speculated the trade-off between TAG and sugars in HME ryegrass leaves alters the leaf sugar homeostasis which in turn mitigates negative feedback on photosynthesis (Beechey-Gradwell et al., 2020; Cooney et al., 2021). However, we observed that HME ryegrass with leaf FA contents above 6.8% DW (TAG > 2.5% DW) appears to have adverse effect on the growth when compared to non-transformant (NT). Understanding the molecular and biochemical changes during TAG accumulation in the non-seed tissues may help to explain these results.

The *de novo* biosynthesis of FA and TAG synthesis places high demands for both energy and reductants which is provided by sugars and the sugars themselves also have multiple roles in signalling pathways including sink regulation of photosynthesis (Xu and Shanklin, 2016; Paul and Foyer, 2001, and references therein). Regulation of photosynthesis requires a communication between sugar sensing and redox chemistry; these can be orchestrated in response to changing environments (Queval and Foyer 2012). Of the environmental factors, light has a substantial effect on photosynthesis over the period of the day. Low light density directly influences photosynthesis and photomorphogenesis, which in turn impacts on the agronomic traits of plants. Reduction of biomass production and higher membrane lipid peroxidation are two common plant responses to adverse low light, especially during winter when this is combined with low temperatures (Gaba and Black, 1983; Höglind et al., 2011). Low irradiance causes limitations of photosynthesis by inhibiting photosynthetic efficiency by lowering levels of PSII, Rubisco, electron transportation rate, and CO_2_ assimilation (Leong and Anderson, 1984). To enhance the tolerance to low lighting, plants improve expression of some specific genes and proteins such as spermidine synthase, ATP synthase β-subunit, and Rubisco activase (Lu et al., 2019, and references therein). It is important to understand how the HME ryegrass perform and acclimate to lower light environment such as those experience under sward (low light field canopy) conditions (Faurie et al., 1996). Does shading affect FA and TAG accumulation in HME leaves? Does elevated photosynthesis and greater biomass created in the HME due to micro-lipid sink overwhelm shading reduced photosynthetic capacity? In this study, therefore, the performance of HME ryegrass grown under a low light density (150-250 µmol m^-2^ s^-1^), was compared to plants grown under standard lighting (600-1000 µmol m^-2^ s^-1^). FA, sugars, growth, photosynthetic parameters and transcriptomic analysis provided evidences into mechanisms fundamental stable lipid storage, sink strength and photosynthesis.

## RESULTS

### Leaf and Sheath FA and Water-Soluble Carbohydrate Levels are Altered in HME Ryegrass and Influenced by Light Levels

Previously generated multiple independent HME lines (HMEs) that co-express DGAT and cys-OLE were screened for their lipid accumulation in the leaf. In this study we selected three of the HMEs (HME30, HME34, and HME42) because these lines represented a range of leaf lipid profiles and total leaf lipid content. Plants (including NT) were grown to undertake a detailed examination of the influence of different photosynthetic irradiance levels on lipid content, growth, photosynthesis and gene expression in the leaves.

Leaf FA profiles of the three HMEs were altered significantly as genotype x light treatment interactions. This included an increase in the proportions of 18:1 and decreases in 16:0, 16:1, and 18:3 (Figure 1). HMEs also have increased proportions of 18:2; 20:0, and 20:1 but these proportions were not affected by the low light treatment. Furthermore, 22:1 was detected in HMEs but not in NT; the level increased in low light. Between the HMEs, HME34 had slightly higher proportions of leaf 18:0, and 22:1. There were small differences in the leaf FA profiles of NT grown in standard compared to low irradiance; this included increases in the proportions of 16:0, 18:1, 18:2 and decreases in 16:1, and 20:1. Ryegrass sheaths are the lower part of subtending leaves, forming a pseudostem, have different FA profile and content to that of the leaf. Compared to NT the sheath FA profiles of HMEs under standard light showed increases in 18:1, 18:2, 20:0, and 22:1 and decreases in 16:0, 16:1, 18:3, and 20:1; while only 16:0, 16:1 and 22:1 were affected in low light. The FA profile of the HME34 sheath was different to the other two HMEs in the proportions of 16:0 and 20:1 under the low light treatment. There were only small changes of 16:0, 18:2 in sheath FA of NT in response to low light treatment.

**Figure 1.**
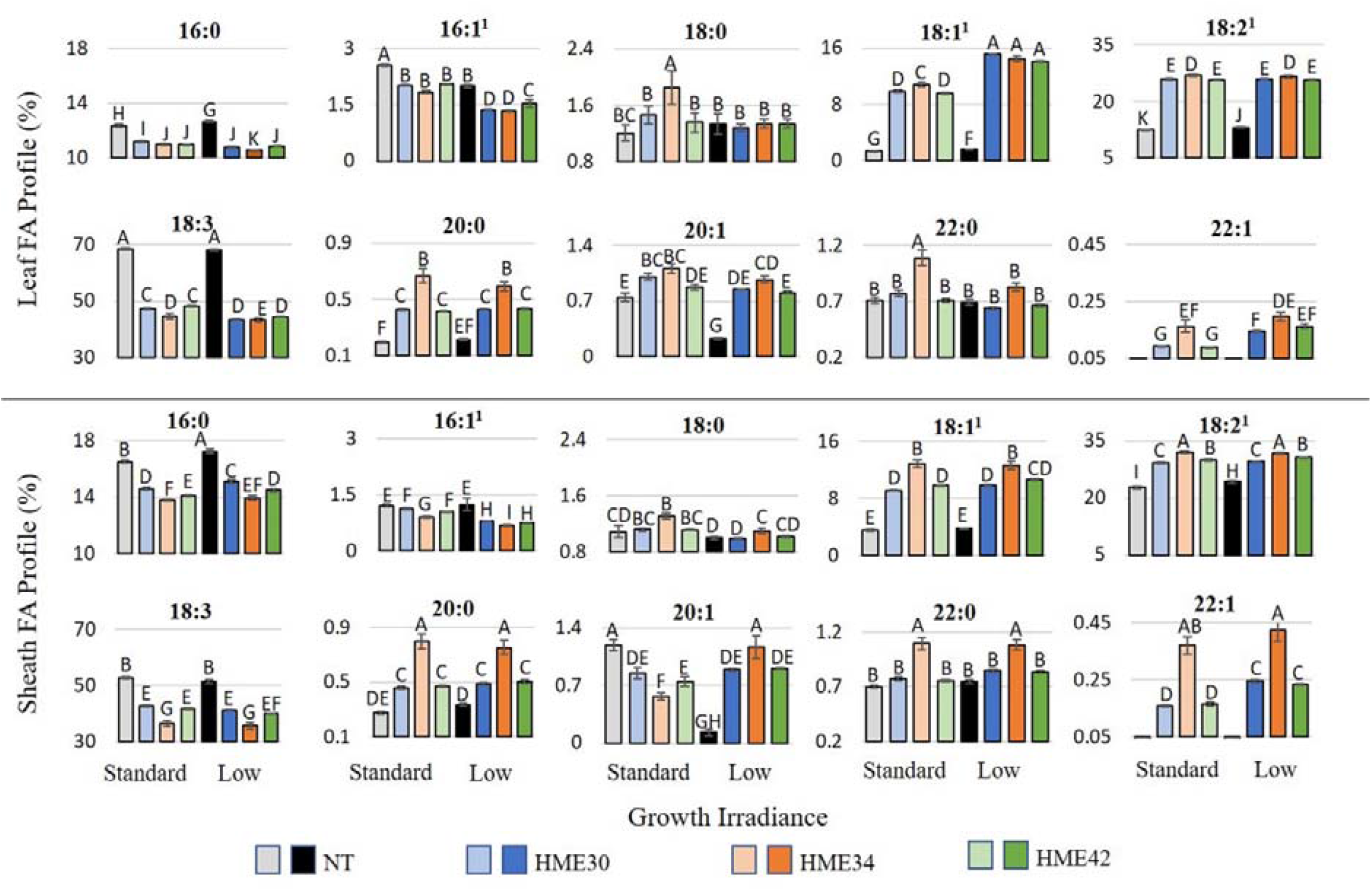
Effect of low light on fatty acid profiles of leaf and sheath total fatty acids extracted from material of high lipid ryegrass and control harvested after 21 days regrowth. Data presented as mean percentages of total fatty acids ±SE from separate analysis (df = 143, *n = 9 or 10*). ^A-K^Alphabets indicated significant differences (*P* < 0.05). ^1^Data were analyzed using log-transformation.

Table 1 shows FA levels for leaves and sheaths harvested at 21 days of regrowth. FA levels accumulated to a higher content in the leaves than in the sheath in all genotypes and under both light treatments. Compared to NT, all three HMEs had significantly elevated leaf FA (∼55-75 %) and sheath FA (∼29-72%) in both light conditions (*P* < 0.001). Low light significantly increased the leaf FA in both NT and HMEs but in sheath only HMEs significantly increased FA. The line with the highest accumulation of leaf lipids (HME34) also had the highest accumulation of cys-OLE in both light conditions (Table 1 and Supplementary Figure S1).

**Table 1.**
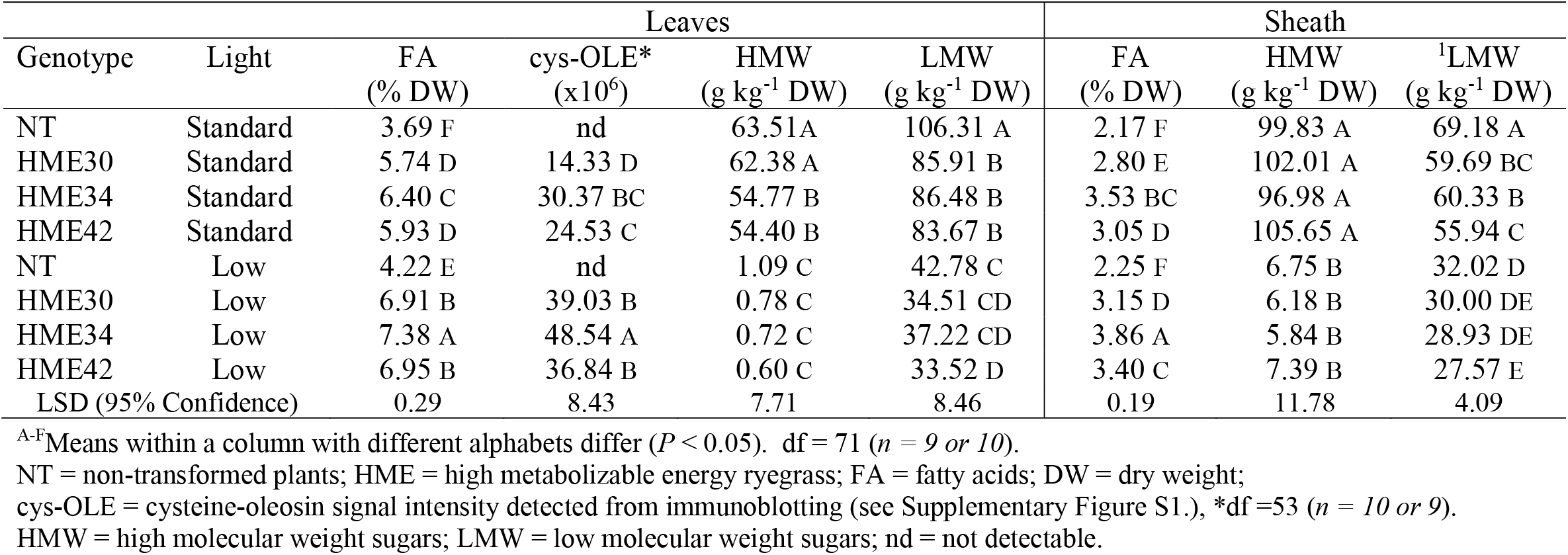
Effect of low light on total fatty acid and sugars in high lipid ryegrass and control.

Both leaf and sheath sugar concentrations decreased significantly when the plants were grown under low light (*P* < 0.001) (Table 1); the decrease was greatest in the high molecular weight (HMW) fraction. HMW-sugars (primarily fructans) accumulated to a higher concentration in the sheath than in the mature leaves; the opposite was true for the low molecular weight (LMW) sugar fraction. All plants had lower levels of leaf and sheath HMW-sugars under low light. HMW-sugars were significantly lower in the leaves of two HMEs (HME34 and HME42) than the NT under standard lighting, however statistical analysis showed no significant difference of leaf and sheath HMW-sugars between genotypes under low light conditions. In both light conditions, HME leaf and sheath LMW-sugars were lower than NT (*P* < 0.001).

### Effect of Low Light on the Growth of HME Ryegrass

All relative growth rates (RGR) of plants grown under standard lighting were significantly greater than plants grown under low light (Figure 2). When comparing plants grown under standard light the overall plant RGR was significantly greater in all HMEs compared to the NT; in descending order the main organs contributing to the elevated RGR were roots, leaves, sheath (Figure 2). In contrast, there was no significant difference in the overall RGR of any line when the plants were grown under low light; this was reflected by the large drop of elevated root RGR trend under this condition.

**Figure 2.**
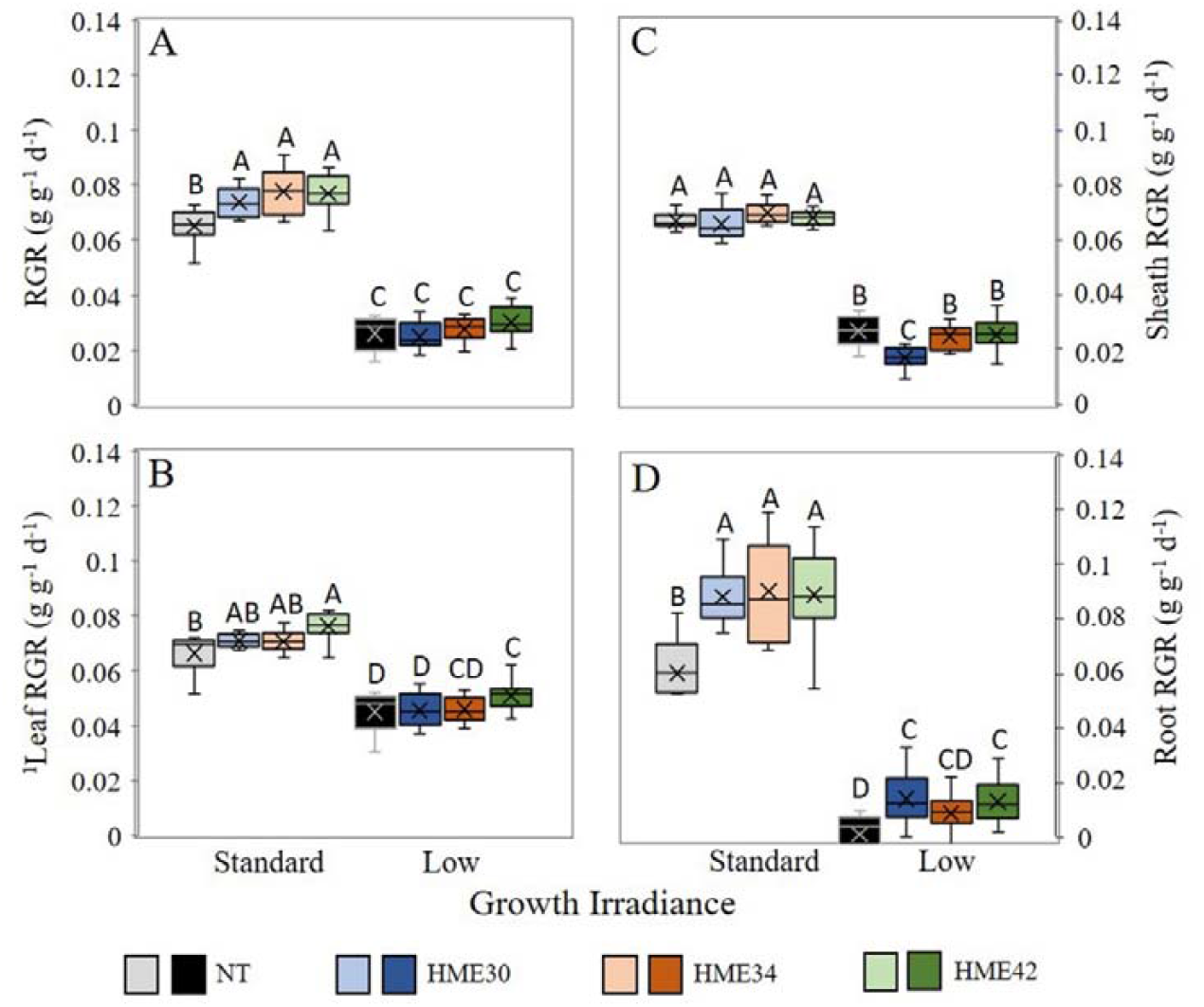
Box and whisker plots of relative growth rate (RGR, g g^-1^ d^-1^) of high lipid ryegrass and control grown under standard and low lights. (A) RGR of total mass (B) leaf RGR (C) sheath RGR and (D) root RGR. ^A-D^Alphabets indicated significant differences (*P* < 0.05). df = 71 (*n = 9 or 10*). ^1^Data were analyzed using log-transformation. WT = non-transformed plants; HME = high metabolizable energy ryegrass; RGR calculated as (ln W_2_ – ln W_1_)/ (t_2_-t_1_), where W_1_ and W_2_ were plant dry weights at times t_1_ and t_2_.

Plants adapt to low light conditions by increasing both leaf area and chlorophyll content in an attempt to capture as much light as possible. In this study we found no significant differences in specific leaf area (SLA) between the HME lines and NT whereas leaves formed under low light having on almost twice the SLA (164-179%) in comparison with standard light leaves (Table 2). Associated with this is an elevation of the shoot to root ratio (L/R) and leaf water content as well as decreases of leaf carbon content (per unit dry matter) and leaf nitrogen content (per unit area, N_area_). There were significant differences in each of these parameters when comparing all plants grown under two different levels of irradiance (Table 2). When comparing between plants grown under low light however, only the leaf carbon content showed a significant difference, where the NT was lower than all HMEs. In comparison, under standard lighting there were significant differences between the HMEs and NT in all these parameters except the leaf nitrogen content. More specifically, under standard light the HMEs have reduced L/R, increased leaf carbon, water and chlorophyll contents. The higher leaf water contents of the HMEs under standard light compared to NT suggest that photosynthesis per unit of water (water use efficiency) in the HMEs may be more efficient; however, it could also reflect the greater ability of the comparatively lower L/R enables them to keep the leaves more turgid. In standard lighting, although three HMEs had higher chlorophyll content (DW basis) than the NT, only one line (HME42) showed significantly higher chlorophyll content per unit leaf area. Under low light, all HMEs and NT exhibited a significant increase in chl per DW, compared to those plants grown under standard light. However, no significant difference in chl was identified between HME and NT at this irradiance. It should be noted however, the influence of light was the opposite when determining the levels of chlorophyll content per unit leaf area.

**Table 2.** Effect of low light on specific leaf area, leaf-to-root ratio, nitrogen content per leaf area, carbon content per dry mass, leaf water, and chlorophyll in high lipid ryegrass and control.

| Genotype | Light | SLA<sup>1</sup><br>m<sup>2</sup> kg<sup>-1</sup> DW | <sup>1</sup>L/R<br>g g<sup>-1</sup> DW | N<sub>area</sub>*<br>g m<sup>-2</sup> | % C* | Leaf water<br>% | Chl<br>mmol kg<sup>-1</sup> DW | Chl<sub>area</sub><br>mmol m<sup>-2</sup> |
| --- | --- | --- | --- | --- | --- | --- | --- | --- |
| NT | Standard | 18.79 B | 0.81 B | 2.32 A | 42.87 B | 71.77 D | 9.59 D | 0.51 BC |
| HME30 | Standard | 19.12 B | 0.59 C | 2.25 A | 44.10 A | 75.47 C | 10.45 C | 0.55 AB |
| HME34 | Standard | 19.83 B | 0.53 C | 2.22 A | 44.36 A | 75.75 C | 10.73 BC | 0.55 AB |
| HME42 | Standard | 19.68 B | 0.59 C | 2.20 A | 44.37 A | 77.15 B | 11.45 B | 0.59 A |
| NT | Low | 32.35 A | 1.85 A | 1.66 B | 41.14 E | 83.90 A | 15.16 A | 0.47 CD |
| HME30 | Low | 34.24 A | 1.73 A | 1.61 B | 42.12 D | 83.92 A | 14.59 A | 0.43 D |
| HME34 | Low | 34.59 A | 1.80 A | 1.60 B | 42.51 C | 83.87 A | 15.14 A | 0.45 D |
| HME42 | Low | 32.20 A | 1.75 A | 1.71 B | 42.32 CD | 83.84 A | 14.76 A | 0.45 CD |
| LSD (95% Confidence) |  | 0.19 | 0.21 | 2.78 | 0.28 | 1.12 | 0.75 | 0.05 |

<sup>A-D</sup>Means within a column with different alphabets differ ( $P < 0.05$ ). df = 71 ( $n = 9$ or $10$ ), \*df = 32 ( $n = 5$ ).
<sup>1</sup>Data were analyzed using log-transformation.
NT = non-transformed plants; HME = high metabolizable energy ryegrass; SLA = specific leaf area (projected leaf area per unit leaf dry mass) L/R = leaf-to-root ratios; N<sub>area</sub> = total leaf nitrogen per unit of leaf area; % C = carbon content per dry mass; Chl = chlorophyll; Chl<sub>area</sub> = leaf chlorophyll per unit of leaf area.

### Elevated Photosynthesis in HME Ryegrass Does Not Occur in Low Light

In agreement with our previous observations (Beechy-Gradwell et al., 2018 & 2020), under standard lighting and compared to NT, the HMEs had elevated CO_2_ assimilation rates (A), transpiration rates (E), stomatal conductance (*g*_*s*_), electron transportation rates (*J*) and quantum efficiency of photosystem II (ϕ_PS2_) (Table 3). However, under low light there were no significant differences in any of these parameters between the HMEs and NT. In all plants the low light treatment significantly reduced A, E, *g*_*s*_, and *J* (55-62%, 49-57%, 47-57%, and 62-64% respectively) while enhancing the ϕ_PS2_ (126-134%) for light harvesting (Table 3). Differences in the response between the A of plants grown in both light densities are associated with a difference in SLA which is a growth measurement that eventually has an effect on photosynthesis.

**Table 3.** Effect of low light on photosynthesis parameters of high lipid ryegrass and control.

| Genotype | Light | A<sup>1</sup><br>μmol m<sup>-2</sup> s<sup>-1</sup> | E<sup>1</sup><br>mmol m<sup>-2</sup> s<sup>-1</sup> | g<sub>s</sub><sup>1</sup><br>mmol m<sup>-2</sup> s<sup>-1</sup> | J<br>μmol m<sup>-2</sup> s<sup>-1</sup> | Φ<sub>PS2</sub> |
| --- | --- | --- | --- | --- | --- | --- |
| NT | Standard | 17.97 C | 1.70 BC | 224.51 B | 148.12 B | 0.50 C |
| HME30 | Standard | 20.87 AB | 1.96 AB | 263.98 AB | 156.92 A | 0.53 B |
| HME34 | Standard | 19.00 BC | 1.79 C | 240.11 B | 153.32 AB | 0.52 BC |
| HME42 | Standard | 22.46 A | 2.21 A | 300.20 A | 158.22 A | 0.54 B |
| NT | Low | 8.00 D | 0.87 D | 119.21 C | 56.30 C | 0.67 A |
| HME30 | Low | 8.52 D | 0.96 D | 133.29 C | 55.60 C | 0.66 A |
| HME34 | Low | 8.73 D | 0.92 D | 125.89 C | 56.98 C | 0.68 A |
| HME42 | Low | 8.54 D | 0.94 D | 128.95 C | 57.35 C | 0.68 A |
| LSD (95% Confidence) |  | 2.09 | 0.28 | 39.40 | 5.65 | 0.02 |

<sup>A-D</sup>Means within a column with different superscripts differ ( $P < 0.05$ ). df = 72 ( $n = 10$ ).670 <sup>1</sup>Data were analyzed using log-transformation.
NT = non-transformed plants; HME = high metabolizable energy ryegrass; A = the rate of photosynthesis per unit leaf area;
E = transpiration rate; g<sub>s</sub> = stomatal conductance; J = electron transportation rate per unit leaf area;673 Φ<sub>PS2</sub> = quantum efficiency of PSII.

### NT and HME Ryegrass RNA-Seq Analysis Under Standard Light

To gain insights into the metabolic process differing between NT and HME ryegrass, we combined RNA sequencing with differential expression analysis. RNA samples were extracted from NT and two HMEs (HME34 and HME42) then processed to determine for differentially expressed genes. Two-well established statistical analysis methods (DESeq and Biostrings) were employed to compare the expression levels. In total 93 up- and down-regulated genes were annotated in this study, calculated as the log_2_ fold-change for the identified genes ranging from ±1 to ±14, and are presented as a heatmap (Figure 3). Within the heatmap we have grouped the genes into related biological processes including: lipid metabolism; light capture and photosynthesis; carbohydrates and the citric acid cycle; sugar signalling and redox maintenance; and nitrogen metabolism and plant and growth development. Each grouping is described in more detail below.

**Figure 3.**
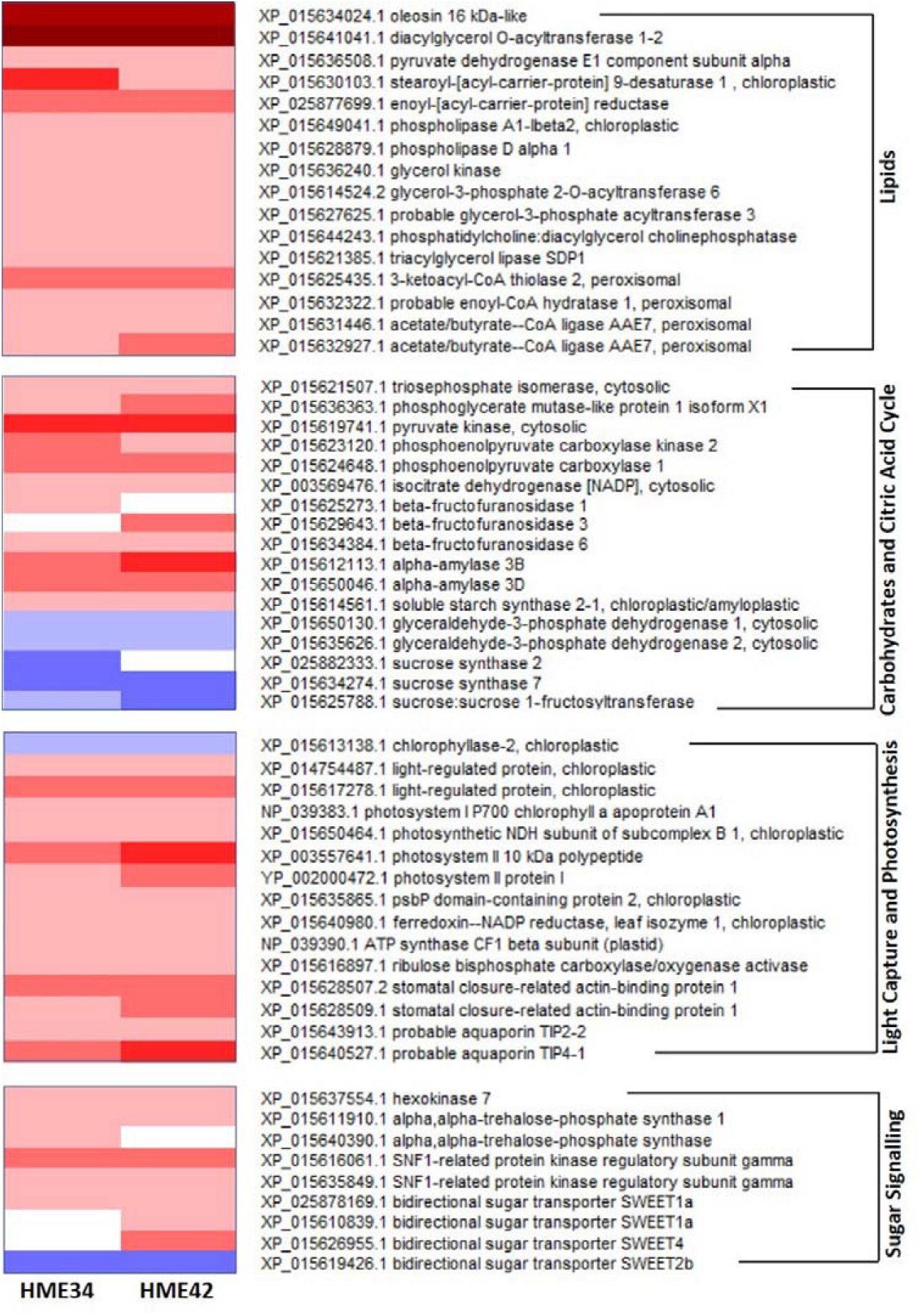
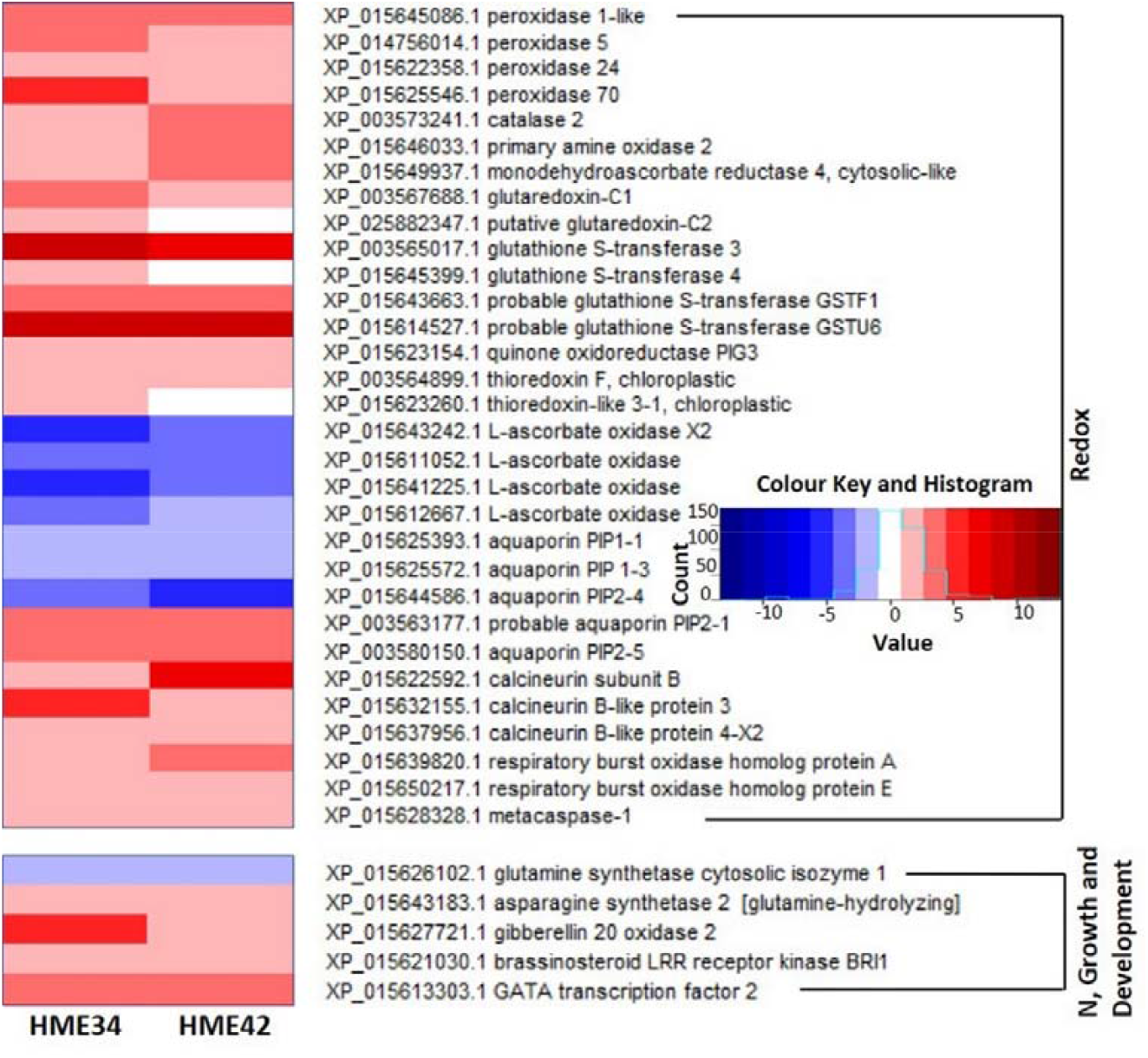
Differential gene expression in high metabolizable energy (HME) ryegrass to non-transformed plants (NT). Colour of the boxes present (Log2)-fold change values relative to NT at significant *P <* 0.001 (*n = 3*).

#### Lipid metabolism

As expected, the recombinant *cys-OLE* and *DGAT* transcripts were detected at relatively very high levels in HME leaves and were not detected in NT. The native genes involved in lipid metabolism that were differentially regulated in HMEs compared to NT were all upregulated; this included genes involved in *de novo* FA synthesis, FA desaturation and elongation, glycerolipid synthesis, TAG catabolism, phospholipid hydrolysis and peroxisomal β-oxidation.

#### Carbohydrates and Citric Acid Cycle

Genes involved in carbohydrate metabolism and the citric acid cycle were both up and down regulated in HMEs compared to NT. Those that were upregulated are involved in glycolysis, the TCA cycle, gluconeogenesis and degradation of carbohydrates in the cytoplasm, as well as hydrolysis of sucrose and starch synthesis in the chloroplast. While those that were downregulated are involved in glycolysis, sucrose and fructan synthesis (Figure 3).

#### Light Harvesting and Photosynthesis

Like the lipid metabolism genes, all genes involved in photosynthesis (with the exception of chloroplastic chlorophyllase which is involved in chlorophyll breakdown) that we identified as being differentially expressed in HMEs were upregulated compared to NT. This included genes directly involved in light harvesting, the photosynthetic electron transport chain and the Calvin cycle (Figure 3). Other genes indirectly influencing photosynthesis by regulating gas, water and solute transport were also upregulated in HMEs.

#### Sugar Signalling Network and Redox Potential

The majority of differentially related genes involved in hexokinase and the sucrose non-fermenting 1-related protein kinase/trehalose 6-phosphate (SnRK1/T6P) signalling pathways were upregulated in HMEs, however, one specific bidirectional sugar transporter was down regulated (Figure 3). Similarly, the differentially expressed genes involved in redox potential were mostly upregulated in HMEs compared to NT. Notable exceptions were L-ascorbate oxidase and several aquaporin plasma membrane intrinsic proteins (*PIPs*).

#### Nitrogen Metabolism and Plant Growth and Development

Two important components of nitrogen assimilation pathway were repressed in HMEs, while two genes involved in gibberellin biosynthesis and brassinosteroid signalling pathways were upregulated (Figure 3).

### Differential Gene Transcriptions in HME Ryegrass to NT in Response to Low Irradiance

In addition to the results shown in Figure 3, Figure 4 shows the normalized transcripts that were influenced by modifying the photon flux in two HMEs (HME34 and HME42) and NT. Although the *cys-OLE* (under the control of *CAB* promoter) was the most abundant transcript we detected under both light treatments its level increased significantly in both HMEs in response to low light (Figure 4A); this agrees with the increased levels of the protein (Table 1). In comparison the *T. majus DGAT1* transcript was only significant different (*P* < 0.001) between HMEs and NT as opposed to different light conditions.

**Figure 4.**
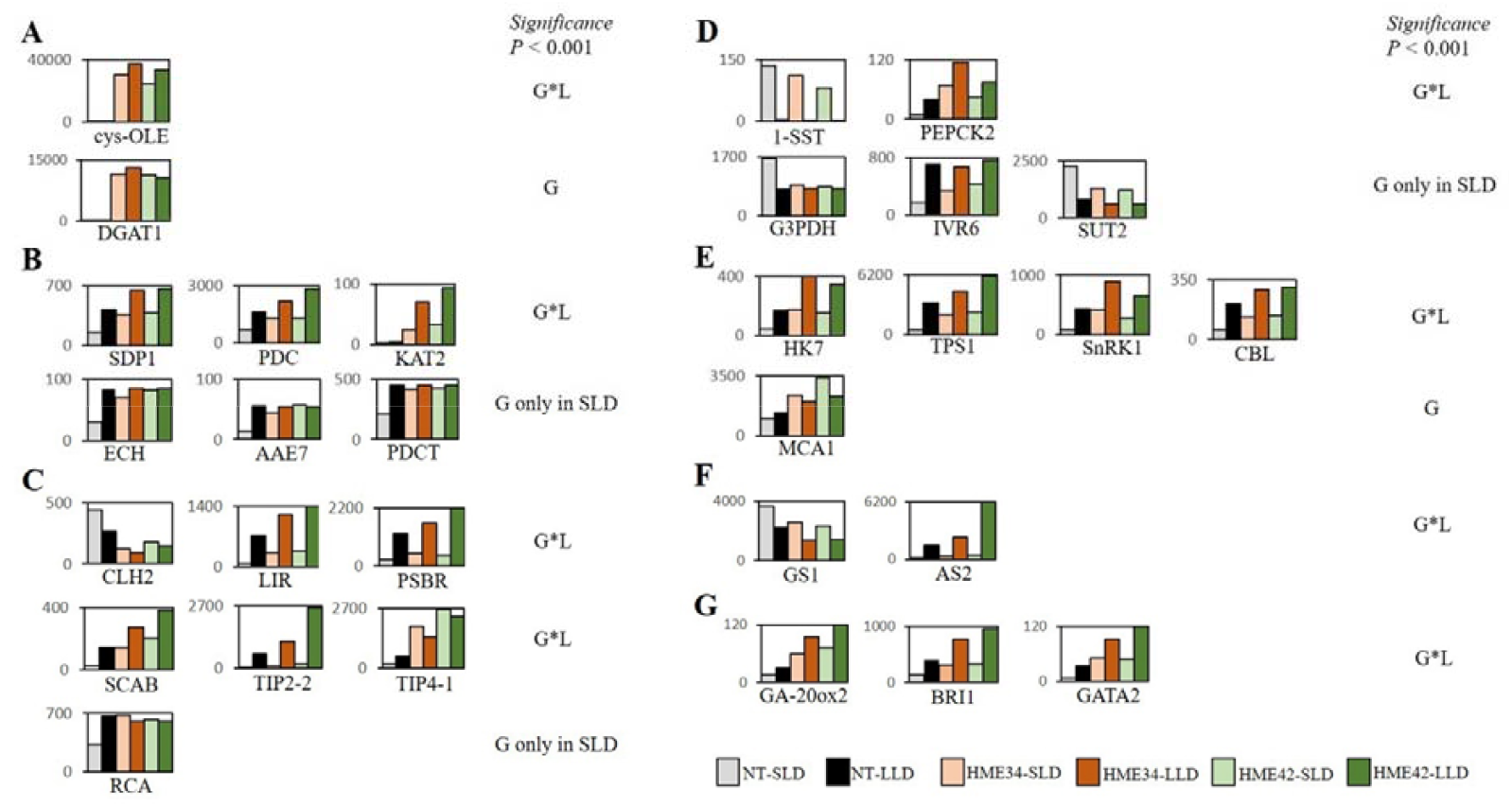
Normalized transcript read counts of annotated genes involving lipid (A and B), light and photosynthesis (C), carbohydrate (D), sugar and oxidative stress signalling (E), nitrogen assimilation (F), and plant growth and development pathways (G) showed differential expression between high metabolizable energy (HME) and non-transformed (NT) ryegrass grown under standard light density (SLD) and low light density (LLD). Each data point represented an average from 3 biological replicates statistically analysed with significance *P* < 0.001 between genotype (G), genotype times light interaction (G*L), and G only in SLD. Gene abbreviation included cys-OLE = cysteine-oleosin; DGAT1 = diacylglycerol acyltransferase 1; SDP1 = triacylglycerol lipase sugar-dependent 1; PDC = pyruvate dehydrogenase E1 complex; KAT2 = peroxisomal 3-ketoacyl-CoA thiolase 2; ECH = peroxisomal enoyl-CoA hydratase 1; AAE7 = peroxisomal acetate/butyrate-CoA ligase; PDCT = phosphatidylcholine:diacylglycerol choline phosphatase; CLH2 = chloroplastic chlorophyllase 2; LIR = chloroplastic light-regulated protein; PSBR = photosystem II 10 kDa proteins; SCAB = stomatal closure-related actin-binding protein; TIP2-2&4-1 = aquaporin tonoplast intrinsic proteins 2-2&4-1; RCA = ribulose bisphosphate carboxylase/oxygenase activase; 1-SST = sucrose-sucrose 1-fructosyltransferase; PEPCK2 = phosphoenolpyruvate carboxylate kinase 2; G3PDH = glyceraldehyde 3-phosphate dehydrogenase; IVR6 = β-fructofuranosidase (insoluble invertase) 6; SUT2 = sucrose transport protein 2; HK7 = hexokinase 7; TPS1 = trehalose 6-phosphate synthase 1; SnRK1 = sugar non-fermented 1 related protein kinase; CBL = calcineurin B-like protein 3; MCA1 = metacaspase 1; GS1 = cytosolic glutamine synthetase 1; AS2 = asparagine synthetase 2 (glutamine-hydrolyzing); GA-20ox2 = gibberellin 20 oxidase 2; BRI1 = brassinosteroid LRR receptor kinase; GATA2 = G-type light-responsive element transcription factor.

Co-expression of *cys-OLE* and *DGAT1* led to the upregulation of a number of genes involved in chloroplastic *de novo* FA synthesis and in the cases of triacylglycerol lipase sugar-dependent 1 (*SDP1*), plastidic pyruvate dehydrogenase E1 complex (*PDC*), and peroxisomal 3-ketoacyl-CoA thiolase 2 (*KAT2*) there were further significant increases under low light conditions (Figure 4B). In contrast, peroxisomal enoyl-CoA hydratase 1 (*ECH*), peroxisomal acetate/butyrate-CoA ligase (*AAE7*) and phosphatidylcholine:diacylglycerol choline phosphatase (*PDCT*) were upregulated to approximately the same level in both light conditions whereas the same genes in the NT only matched these levels of expression when the plants were grown under low light conditions (Figure 4B).

Three genes involved in light harvesting genes: chloroplastic chlorophyllase 2 (*CLH2*), chloroplastic light-regulated protein (*LIR*); and PSII 10 kDa proteins (*PSBR*) and three genes involved in regulating water flow: stomatal closure-related actin-binding protein (*SCAB*); aquaporin tonoplast intrinsic proteins 2-2 & 4-1 (*TIP2-2 & 4-1*) all showed significant genotype x light treatment interactions (Figure 4C). Where *CLH2* expression was significantly reduced in the HMEs under both light regimes and compared to the expression under standard lighting there was a significant reduction in NT under low light. *LIR, PSBR, SCAB* and *TIP2-2* all showed significant increases under low light with the greatest changes occurring in the HMEs, while *TIP4-1* was highly expressed in both HMEs under both light conditions in comparison to NT. Rubisco activase (*RCA*) expression on the other hand was significantly reduced only in the NT under standard lighting (Figure 4C).

The expression of three genes involved in sugar metabolism sucrose-sucrose 1-fructosyltransferase (*1-SST*); phosphoenolpyruvate carboxylate kinase 2 (*PEPCK2*); and hexokinase 7 (*HK7*) showed significant genotype x environmental interactions (Fig 4D). Whereas four sugar metabolism related genes showed a significant difference between genotypes only in full light conditions: glyceraldehyde 3-phosphate dehydrogenase (*G3PDH*); sucrose transport protein 2 (*SUT2*); β-fructofuranosidase (insoluble invertase) 6 (*IVR6*); and mitochondrial malate dehydrogenase (*MDH*). Three genes related to sugar homeostasis and SnRK1/T6P signalling had significant interactions between the genotype and light regime: *HK7*; trehalose 6-phosphate synthase 1 (*TPS1*); and *SnRK1* (Figure 4E). A similar response was observed for calcineurin B-like (*CBL*), a gene involved in Ca^2+^ signalling related to oxidative stress (Figure 4E). Metacaspase (*MCA1*) showed a significant genotype difference where the expression level was higher in both HMEs. It should be noted however, that although not significant, the expression level was highest in the HMEs in standard light conditions whereas in NT the opposite was true (Figure 4E).

In addition, the expression of two genes involved in nitrogen assimilation: glutamine synthetase 1 (*GS1*) and asparagine synthetase (*AS2*) and three genes involved in regulating plant growth and development: GA-20 oxidase 2 (*GA-20ox2*), brassinosteroid LRR receptor kinase (*BRI1*), and *GATA* transcription factor 2 all showed significant genotype x light treatment interactions (Figures 4F and 4G). The expression of *AS2, GA-20ox2*, and *BRI1* were all upregulated in both HMEs under standard lighting and there was a further significant increase under low light conditions whereas the opposite was true for *GS1*.

## DISCUSSION

In this study, the performance of HME ryegrass grown under different irradiance levels was investigated. FA, sugars, growth, photosynthesis parameters and transcriptomes were analysed. Overall results suggested molecular mechanisms involve carbon partitioning between TAG accumulation and sugar signalling, and redox potential were differentially regulated in the HME plants compared to the NT. Further discussions related to these altered mechanisms are as follows.

### Lipid Metabolism and the Effect of Low Light

Overexpression of DGAT and cys-OLE promoted carbon flux to FA and TAG synthesis and resulted in changes to the expression pattern of several gene transcripts involving FA elongation, FA desaturation and TAG formation. Increased transcript of a plastidic *PDC* in the HMEs suggests the plants favoured *de novo* FA synthesis in the chloroplasts by contributing to acetyl-CoA and NADH production from pyruvate (Ke et al., 2000). *PDC* transcripts of NT and HMEs were increased in response to low irradiance and was concomitant with a higher FA content of both genotypes. Upregulation of the chloroplastic *stearoyl-ACP Δ9-desaturase* transcript in the HMEs correlates with the higher proportion of 18:1 detected in the HME plants (Winichayakul et al., 2020).

Leaf FA composition varies with environmental factors, such as temperature and light and under low irradiance, the light harvesting capacity is usually increased by enhancing the chlorophyll concentration and membrane lipid content of thylakoid per granum (Lichtenthaler et al., 1981; Fuhrmann et al., 2009). Plastid lipid biosynthesis and the degree of thylakoid stacking were significant higher in *Arabidopsis* grown under low light than the plants grown under high light (Yu et al., 2021). Increased leaf FA and change in FA profiles of NT and HMEs grown under low light likely reflect the altered chloroplast ultrastructure of plants that had adapted to this irradiance (Melis, 1984). It has to be noted that some of the change in leaf FA composition and cys-OLE protein of HMEs under low light may also be influenced by the Rubisco small subunit and *CAB* promoters used to control the expression of *DGAT1* and *cys-OLE*, respectively. Several studies in both monocot and dicot species have shown that the *Rubisco small subunit* and *CAB* promoter activities are positively regulated by light with specifically expressing in chloroplast-containing cells at their highest levels during early leaf development (reviewed in Chory and Susek, 1994; Feng et al., 2014; Mukherjee et al., 2015 and references therein). Under low light, although activity of the *CAB* promoter used to drive the expression of *cys-OLE* was expected to be repressed, low light HME leaves contained higher cys-OLE than high light leaves. It is remarkable how plants fine-tune FA and glycerolipid biosynthesis to their cellular metabolic need in response to long-term changes in light conditions.

### Sequestering of Carbon Forms under Different Irradiances

Plants fix inorganic CO_2_ by carboxylating ribulose-bisphosphate, producing a phosphorylated intermediate which is subsequently reduced to triose phosphates (TP) for translocation to the cytosol primarily, but not exclusively, for starch and sucrose synthesis (McClain and Sharkey, 2019). Upregulation of cytosolic *triosephosphate isomerase, phosphoglycerate mutase*, and *pyruvate kinase* gene transcripts in HMEs suggests there have been an increase of carbon flow from phosphorylated intermediates into pyruvate. It has been shown that increased lipid biosynthesis in vegetative tissues can lead to a higher capacity of TP utilization; although synthesis of lipid from pyruvate accounts for only 1-3% of fixed carbon (Bao et al., 2000; Sanjaya et al, 2011). Higher TP utilization can also result in inhibition of starch and sucrose synthesis (McClain and Sharkey, 2019). In our standard lighting environment, we observed downregulation of the leaf cytosolic *sucrose synthase* and *1-SST* gene transcripts in the HMEs which correlates with the lower levels of leaf LMW-sugars and fructans, respectively. *L. perenne* stores very little starch; most of the photosynthetically fixed carbon is directed into sucrose and fructan synthesis (Pollock & Cairns 1991; Fischer et al., 1997). It is the fact that low light limits carbon capture of the plants and reduces sugar available for transportation through the plant cells. As such biosynthesis of fructan by 1-SST and storage of high molecular weight carbohydrates, which usually occurs when carbon assimilation exceeds carbon use, is therefore significantly reduced (Pollock & Cairns 1991).

### Regulation of Sugar Signalling in Response to Leaf Lipid Sink

Sugars play a role in the feedback inhibition of photosynthesis though hexokinase and SnRK1/T6P signalling pathways; the processes which have been well-known as the primary components of sugar homeostasis in plants (Rolland et al., 2002; Li and Sheen 2016). When the HME plants were grown in standard lighting conditions they showed increases in the expression of these three sugar sensing regulators. HK7 is a multifunctional hexokinase, regulating glycolytic flux in the cytosol and acting as a glucose sensor in the nucleus (Kim et al., 2016; Aguilera-Alvarado and Sánchez-Nieto 2017). As such it appears that HK7 activity is elevated in the HMEs to provide more substrate for the oxidative pentose phosphate pathway and subsequent NADPH production which is required for *de novo* FA synthesis. SnRK1 is a central regulator in adjusting cellular energy and metabolism during starvation, stress and growth-promoting conditions (Tsai and Gazzarrini 2014; Wurzinger et al., 2018). As such, the upregulation of SnRK1 in the HMEs is unsurprising given we have shifted the level of energy and the form of energy. Expression of genes involved in photosynthesis were shown to be activated by SnRK1 overexpression although there is limited evidence for a direct link of this signalling pathway (Baena-Gonzalez et al., 2007; Cho et al., 2012). Antagonistically to SnRK1 activity, T6P elevation by overexpressing TPS1, an enzyme involved in T6P synthesis, can positively impacted photosynthesis (Pellny et al., 2004). However, rather than have a direct sink effect on photosynthesis, T6P changes the distribution of photoassimilate in phloem cells through *SWEET* genes, triggering sugar demand within the photosynthesis cells (Oszvald et al., 2018). The altered expression of some *SWEET* genes and TPS1 in the HMEs suggests a role for T6P in balancing sugar homeostasis between source and sink. It is also likely that the increased growth of HMEs results in a greater demand for sugar in the sink tissue which would also reduce the feedback signals that normally downregulate photosynthesis allowing a greater carbon capture (Paul and Foyer 2001; Paul and Eastmond 2020; Beechey-Gradwell et al., 2020).

### Light Harvesting and Photosynthesis of the HME and the Effect of Low Light

Under standard lighting, the HMEs enhance their capacity for light absorption by upregulating light regulating proteins as well as downregulating the chloroplastic CLH2, an enzyme involved in chlorophyll breakdown. Upregulation of PSI-related, PSII-related and *RCA* transcripts, including two key photosynthesis regulators; chloroplast ATP synthase and ferredoxin-NADP reductase (FNR), in the HMEs grown under standard light support the greater photosynthetic activity of the plants. Elevated chloroplast *ATP synthase* and *FNR* transcripts suggests an increase of both linear and cyclic electron flow around photosystem I. This mechanism is essential for photosynthesis, in particularly the cyclic electron transfer that produces ATP required for driving CO_2_ fixation (Mulo 2011 and references therein; Bhaduri et. al., 2020). In addition, an increased RCA transcript in the HMEs associated with the plants’ greater electron transport suggests modulation of the Rubisco activity may occur (Perdomo et al., 2017).

In low light, both NT and HMEs enhanced their capture of light energy by increasing chlorophyll and SLA; although this led to a lower Chl_area_ it improves light transmission allowing higher ϕ_PS2_ and potentially increased photosynthesis by the lower leaves (Praba et al., 2011; Hamblin et al., 2014). Low light negatively affects *J*, E and particularly *g*_s_ resulting in higher intercellular CO_2_ and increased leaf H_2_O which limits diffusion capacity and therefore constrains CO_2_ assimilation (Liu et al., 2014; Zhu et al., 2017). Various genes related to gas, water and solute transport such as *SCAB* and *TIP2-2, TIP4-1* were upregulated in the HMEs, indicating change associated with stomatal conductance and vascular functions, respectively. These three gene transcripts were also increased in NT and HMEs in response to low light. SCAB controls guard cell movement and its inhibition interrupts stomal closure and opening (Zhao et al., 2011). Moreover, it has been suggested that SCAB plays an important role in plant growth under changes of light quality by regulating actin filament dynamic (Wang et al., 2015). Given the abundance of TIP aquaporins in the plant tonoplasts and their redundant roles in membrane transport of water and other small molecules, an accurate explanation for the upregulation of TIP2-2 and TIP4-1 transcripts in the HMEs may be confound without further investigation (Maurel et al., 2015).

### Changes in Redox Potential in Response to Leaf Lipid Sink

The HMEs also differentially manipulate both enzymatic and non-enzymatic mechanisms involved in redox homeostasis such as peroxidases, catalase, amine oxidase, and ascorbate-glutathione cycle components as well as L-aspartate oxidase. Combined, these function to remove conjugated glutathione, detoxify peroxidised lipids from the cells and transport oxidative molecules between the cells (Foyer and Noctor 2011). Ascorbate oxidase has a prominent role in the redox homeostasis; therefore, it may be necessary for the HMEs to manipulate the level of this enzyme in order to maintain the redox balance. In addition, increased levels of ascorbate may reduce the recycle of ascorbate from oxidised forms (dehydroascorbate and monodehydroascorbate) and in doing so reserve more reduced forms of glutathione and NADPH (Asada, 2006). Further experiments to determine the change in redox potential of the HME plants will be required.

HMEs appear to differentially regulate specific PIP aquaporins. Several researches have explored the role of PIP to facilitate the retrograde signalling of oxidative compounds, particularly H_2_O_2_ (Bienert and Chaumont 2014). A few possible contributors that may lead to the accumulation of H_2_O_2_ and the altered redox regulation in the HMEs are the elevated levels of FA β-oxidation in the peroxisomes and the greater electron transport and photosynthesis (Niu and Liao, 2016). In this study we found that numerous transcripts (e.g. *SDP1* lipase, *KAT2*) involving TAG catabolism and peroxisomal oxidation of FA are increased in the HMEs; similar findings were reported in high oil tobacco (Vanhercke et al., 2017). The functions of SDP1 and other TAG lipases are to synergistically direct FA toward β-oxidation, thereby maintaining membrane lipid homeostasis (Fan et al., 2014). KAT2, an enzyme involved in the final step of β-oxidation of FA, was shown to regulate the phytohormone ABA signalling partly through modulating reactive oxygen species (ROS) homeostasis in plant cells (Jiang et al., 2011). Both *SDP1* and *KAT2* were also upregulated in NT and HMEs in response to low light with significant genotype x light interaction.

ROS such as H_2_O_2_ are not only considered as cell destructive compounds but act as a signalling molecule in various metabolic processes, including Ca^2+^ signalling and programme cell-death (Steinhorst and Kudla, 2013; Coll et al., 2010; Černý et al., 2018). Both HMEs had upregulated levels of transcripts for CBL, respiratory burst NADPH oxidase, and MCA1. CBL also increased in NT and HMEs in response to low light while MCA1 only increased in HMEs with no significant difference due to light conditionings. A direct interconnection between CBL-mediated Ca^2+^ and ROS signalling in plants was reported as evidence for a synergistic activation of the NADPH oxidase, the key producer of ROS in plants (Drerup et al., 2013). Plant metacaspases, represent a family of cysteine proteases which are structurally related to metazoan caspases but are different in their substrate specificity as they cleave at the arginine and lysine residues (Vercammen et al., 2004). MCA1, a type I metacaspase, was found to be a positive regulator of oxidative stress and developmental-induced cell death (Tsiatsiani et al., 2011). Interestingly, proteome analysis has revealed MCA1 is involved in the dissipation of protein aggregates (Lee et al., 2010). HME may have upregulated MCA1 in response to a cluster of cross-linked cys-OLEs at the closed proximity to the ER during lipid encapsulation. We previously demonstrated that cys-OLE appears to be highly resistant to cysteine protease degradation (Winichayakul et al., 2013) and it would be interesting to examine the stability in the presence of MCA1.

### Regulation of Nitrogen Assimilation and Plant Growth and Development in HMEs and the Effect of Low Light

Our analysis has shown that the leaf N_area_ was not significantly different between the HMEs and the NT but decreased under low light. The expression level of two important components of the nitrogen assimilation pathway (GS1 and AS2) were strongly influenced by both genotype and environment, where the cytosolic *GS1* transcript was repressed in HMEs and further reduced in low light; the opposite was true for *AS2*. Expression of *AS* genes are repressed by sugars and light in a number of plant species (Silvente et al., 2008, and references therein). It has been postulated that the evolutionarily conserved *cis*-regulatory elements of the *AS* promoters mediate sucrose-repression and hexokinase may be involved in this regulation (Irving et al.,2001; Winichayakul et al., 2004). We speculate that the upregulation of *AS2* transcript in HMEs may be due to the plants lower sugar content. As for the plants grown under low light, the *AS2* gene expression in NT and HMEs may be upregulated directly by physical light signalling and/or indirectly by light levels influencing the sugar content of the plant cells. Three distinct cytosolic GS isoforms from Poaceae family have been reported; these isoenzymes are essential for nitrogen remobilization from senescing leaves to fill grain in annual species (Hirel et al., 2001; Obara et al., 2004; Bernard et al., 2008). Regulation of cytosolic GS1 in ryegrass may be different as the plant is a perennial and the plant materials are vegetative tissues. In addition, the upregulation of *AS2* or low sugar may be a feedback signal to reduce cytosolic GS1.

Three genes involved in regulating plant growth and development: *GA-20ox2, BRI1* and *GATA2* are upregulated in HMEs and in low light. Reducing light intensity was shown to result in an inversely proportional increase in gibberellin content which positively correlated with stem elongation (García-Martinez and Gil, 2002). BRI1 is involved in brassinosteroid signalling cascades during plant development, promotion of cell division and elongation. It has been shown that light and brassinosteroid signalling antagonistically regulate the plant developmental switch from skotomorphogenesis to photomorphogenesis through a key light responsive element G-type GATA2 transcription factor (Luo et al., 2010).

## CONCLUSION

In our previous study we reported that creating stable micro-sinks (lipid droplets) within the photosynthetic cell of the HME plants reduces the availability of primary photosynthate (sugars). In turn this reduces the diurnal build-up of carbohydrates and as such prevents the feedback inhibition on photosynthesis. This is the first study to report on transcript expression in these plants. Many of the upregulated transcripts were from genes involved in lipid metabolism, light capturing, and photosynthesis. These were also shown to be modulated by the low intensity of the photosynthetic light. The results from this study suggest that our development of HME ryegrass should provide a promising towards delivering pasture sward (low light field canopy) with higher lipid. However, the secondary effect of HME expression that increases photosynthesis and growth yield of the plants may be not efficiently translated from the laboratory spaced pots to sward canopy, due to shading conditions.

In addition, we also reported that expression of genes involved in sugar signalling and redox homoeostasis was influenced by the accumulation of lipid droplets in the leaf. Further studies on the regulatory network such as nitrogen metabolism, phytohormone signalling, sugar regulation and redox homeostasis are needed to provide an insight into underlying mechanisms which have been altered in response to storing carbon as leaf lipid sink.

## MATERIALS AND METHODS

### Plant Material and Experiment Design

Three independent T_0_ HME transgenic ryegrass lines (HME30, HME34, and HME42) and a respective NT control (Impact cultivar clone 93) were propagated into 1.3 L potting mix soil to generate multiple isogenic clones of each. These transgenic lines were selected due to their range of leaf FA content. Thirty plantlets of 5-tiller each genotype were transplanted into 1.3 L washed coarse sand (Daltons, NZ) flushed with 2 mM KNO_3_ containing nutrient solution (Andrews et al., 1992). Plants were grown in a controlled environment room (22/15 °C day/night temperature, 65-70% humidity, 10 h day-light intensity of 300-400 µmol m ^-2^ s^-1^) and fed with 50 mL of 2 mM KNO_3_ containing nutrient solution, 3 times a week. Pots were rotated along the bench every 2-3 d to provide uniform exposure to the growth conditions.

Three weeks after establishment, randomly selected 10 plants from each genotype were harvested. The remaining 20 plants of each genotype were divided into 2 groups for a comparison between standard (600-1000 µmol m ^-2^ s^-1^) and low (150-250 µmol m ^-2^ s^-1^) light densities. Plants were grown for another 3 weeks and fed with 50 mL of 4 mM NH_4_NO_3_ containing nutrient solution, 3 times a week and every day during the last week. One week prior to the final harvest, plants were measured for photosynthetic parameters.

### Harvest, Leaf Water Content and Relative Growth Rate

Plants were harvested after 3-week establishment and divided into ‘leaf’ (leaf blade and leaf sheath from 6 cm above the sand surface), ‘sheath’ (leaf sheaths on a tiller or pseudostem, 6 cm from the sand surface), and roots (cleaned). These plant materials were oven dried at 65 °C for approximately 3-4 d and weighted for the DW_establish_. At final harvest, plant materials were sampled as above. Small aliquots of leaf were subsampled, weighted for fresh weight (FW^1^), and oven dried for dry weight (DW^1^). The remaining leaf material, sheath and roots were freeze-dried for 3-4 d for DW_final_. Leaf water content was calculated using a classical assessment based on the weight change between fresh and dried leaves as the following:

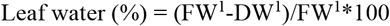

In this study, RGR is preferred than DW in order to eliminate ambiguous differences of DW data resulted from propagation. RGR was calculated from the average of DW_establish_ and individual DW_final_ as the following:

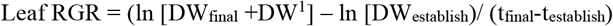

where t is time between established and final harvests.

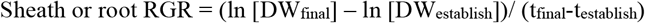

### Lipid and Sugar Analyzes

The lyophilized ground leaves and sheath were analyzed for FA and sugars. FA were extracted from approximately 10 mg DW and methylated according to Browse et al., (1986). FA were verified and analyzed by gas chromatography-flame ionization detector (Shimadzu QP2010, Zebron™ ZB-FAME column (7FD-G033-05)) using Supelco® 37 component FAME standard mix (Merck, CRM47885). The internal pentadecyclic acid (15:0, added prior to methylation) and methylated margaric acid (17:0) standards were used for quantification. Total FA concentration presented as percentage of plant dry weight are the sum of palmitic acid (16:0), palmitoleic acid (16:1), stearic acid (18:0), linoleic acid (18:1), linolenic acid (18:3), arachidic acid (20:0), gondoic acid (20:1), behenic acid (22:0), and erucic acid (22:1).

For water-soluble sugars, approximately a 25 mg sample was extracted in 80% ethanol (v/v) and H_2_O for LMW- and HMW-sugars, respectively (Winichayakul et al., 2020). The colorimetric reaction was developed using anthrone assay as described in Jermyn, (1975) and measured at 620 nm. Sugars were quantified using a dilution series of sucrose and inulin standards.

### Leaf Carbon and Nitrogen Content

The lyophilized ground leaf materials were placed in a drying oven at 30 °C for 15-20 min and approximately 100 mg DW were taken. Samples were repeatedly ground through a 1 mm sieve and analysed for total N and C content by thermal conductivity detection using a combustion elemental analyser (vario MAX cube CNS; Elementar Analysensteme GmbH, Langenselbold, Germany).

### SDS-PAGE Analysis of cys-OLE

Protein samples were prepared by vigorously mixing 10 mg of lyophilized ground material with 150 µL of sterile H_2_O, 200 µL of 2X loading buffer (1:2 diluted 4X lithium dodecyl sulfate sample buffer [Thermo Fisher NP0007], 8 M urea, 4% [v/v] triton X-100, and 5% [v/v] β-mercaptoethanol), and 40 µL of 10X reducing agent (Thermo Fisher NP0004). Equal volumes of leaf protein were separated by SDS-PAGE (Mini-PROTEAN® TGX stain-free™ precast gels; Bio-Rad system) and cys-OLE was detected by immunoblotting as described in Winichayakul et al., (2013). Chemiluminescent activity was developed using WesternBright ECL spray (Advansta, CA) and visualized by the Bio-Rad ChemiDoc™ imaging system. Volume intensity of protein bands was quantified using Image Lab™ software for PC version 5.2.1 (Bio-Rad).

### Chlorophyll Concentration

Chlorophyll (*Chl*) was extracted from approximately 15 mg of lyophilized ground leaf materials with 3 mL of 95% [v/v] ethanol. Extraction tubes (sealed with Teflon lined screw cap) were kept at room temperature in dark for 4 h with interval thoroughly mixing. The extracts were 1/4 diluted before colorimetric measurement. Concentration of chlorophyll in extracts was measured spectrophotometrically (VersaMax™ microplate reader) at the wavelengths of 648 nm and 664 nm and calculated for *Chl*_*a*_ and *Chl*_*b*_ contents as described by Miazek and Ledakowicz (2013). Microplate absorbance from 200 µl of an ethanol extract was adjusted to 1-cm pathlength spectrophotometer with modification of 1/0.51 co-factors according to a report by Warren (2008).

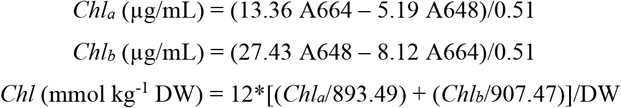

### Photosynthetic Gas Exchange

Three tillers were selected per plant, and on the youngest fully expanded leaves, net photosynthesis per unit leaf area, transpiration, stomatal conductance, electron transportation rate and quantum efficiency of PSII was analyzed using a Licor 6800 infrared gas exchange system (Licor Biosciences Ltd., Nebraska, USA). Leaves were acclimated under growing conditions: 200 and 700 µmol photons m^−2^ s^−1^ red/blue light for low and standard light treatments respectively, 400 ppm CO_2_, 70% relative humidity and 20 °C for 15 min prior to data-logging. The three leaves were then abscised, photographed, oven dried and weighed. Leaf area was calculated using GIMP 2.8.22 (GNU Image Manipulation Program, http://www.gimp.org) and specific leaf area was calculated as a m^2^ of leaf area per kg DW.

### RNA Isolation and Sequencing

Two HMEs (HME34 and HME 42) were tested for transcriptome analysis in this study. Approximately 50 mg of lyophilized ground leaf materials was used for RNA extraction with the Sigma-Aldrich Spectrum Plant Total RNA Kit, according to the manufacturer’s instructions. Total RNA (2-3 µg) was dried down and preserved in Sigma-Aldrich RNAstable® before being couriered to BGI Tech Solutions (Hongkong) Co., Ltd. BGI Tech prepared and processed libraries for 20M PE100 reads using their DNBseq platform.

### Transcriptome Assembly, Expression Analysis and Annotation

The provided reads were assembled using Trinity v2.7.0 with a minimum contig length of 300 bp. Statistics were derived using BWA and Samtools. Differential expression levels were determined in R using DESeq and Biostrings. Differentially expressed contigs were firstly selected for those with *padj <* 0.001, and then those contigs were further refined by selecting for (Log2)-fold change ranged from ±1 to ±14. Local BLAST searches of the *Oryza sativa* IRGSP-1.0 and the *Brachypodium distachyon* databases provided best-hit identification of the differentially expressed contigs. Metabolic gene classes were taken as assigned by the KEGG database (Kanehisa and Goto, 2000) and UniProt Knowledgebase.

### Statistical Analysis

Experiment data were analysed by one-way or two-way ANOVA using the RStudio version 3.6.0 with a model that included fixed effect of ryegrass genotypes (HMEs and NT) and light densities (standard and low lights). In some cases, a log-transformation was applied to the responses for matching the normality assumption of ANOVA. A multiple comparison of treatments such as Bartlett’s test (homogeneity of variances) and Shapiro-Wilk normality test from ANOVA was used to highlight significant among treatment means while *P* values were adjusted by the BH method (Benjamini and Hochberg, 1995) to control the false discovering rate. Means and LSD are reported, and fixed effects declared significant at *P* < 0.05.

## Supplemental Data

The following supplemental material is available.

Supplemental Figure S1.

## ACKNOWLEDGEMENTS

This manuscript was funded by AgResearch Science Prize 2017 and the New Zealand Ministry of Business, Innovation and Employment through the research programme (C10X1603).

---

**Supplementary Figure S1.**
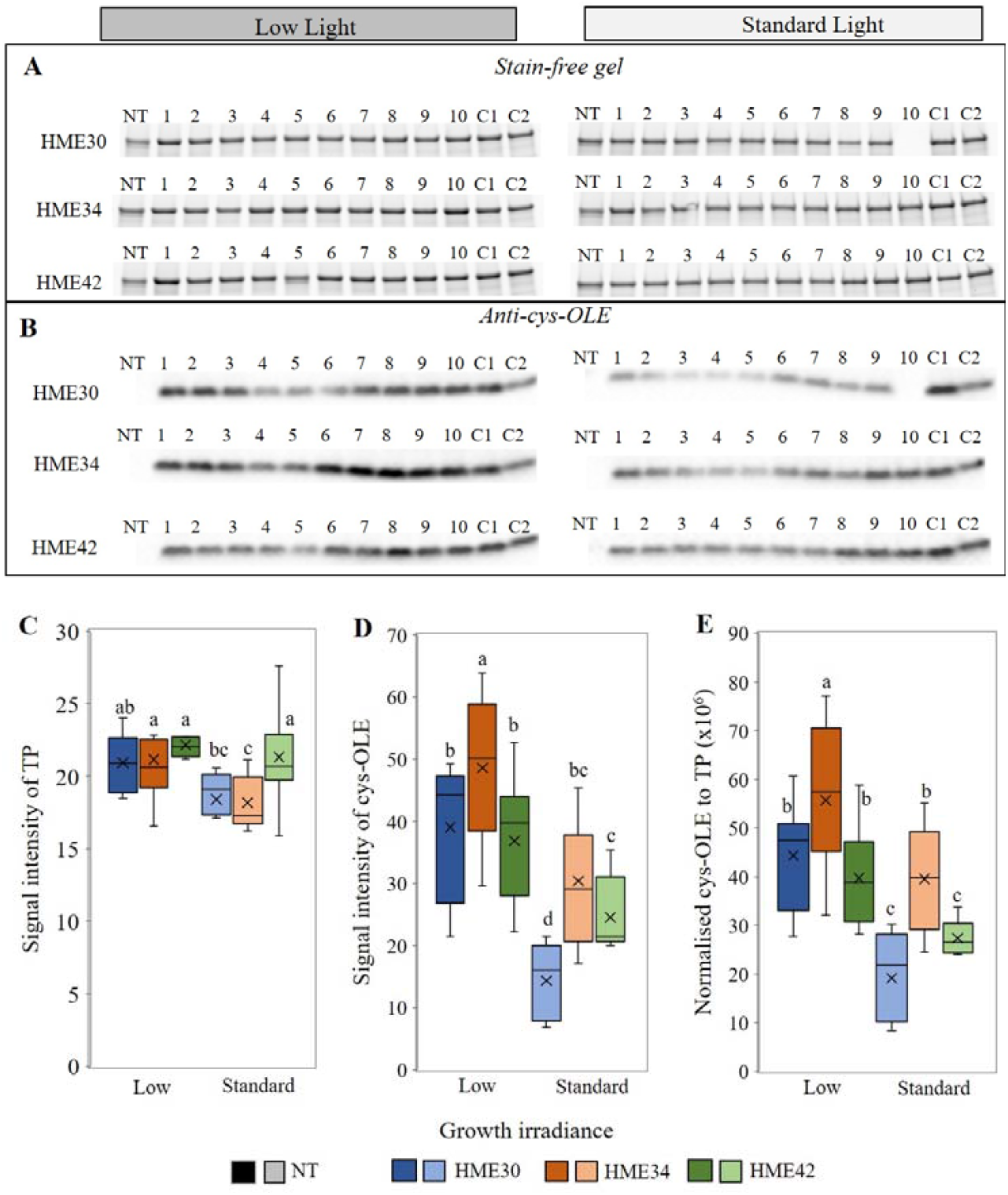
Determination of cysteine-oleosin (cys-OLE) accumulation in leaves of high metabolizable energy (HME) ryegrass and non-transformed plants (NT) grown under low and standard lights. Stain-free gel analysis **(A)** of total proteins extracted from 25 mg of leaf dry material (images showed ∼56 kDa band intensity), and **(B)** immunoblotting results using anti-oleosin antibody; C1 and C2 were controls of protein loading and immunoblotting. **(C)** Box and Whisker plots of signal intensity of total proteins, **(D)** cys-OLE and **(E)** normalised signal intensity of cys-OLE to total proteins (TP). ^a-d^Alphabets indicated statistical difference (*P* < 0.05). df = 53.

